# Critical issues found in “Dissecting cell identity via network inference and in silico gene perturbation”

**DOI:** 10.1101/2024.10.16.618746

**Authors:** Jalil Nourisa, Antoine Passemiers, Sven Tomforde

## Abstract

In the 2023 Nature publication “Dissecting cell identity via network inference and in silico gene perturbation” [1], the authors introduced CellOracle (CO), a novel method leveraging mRNA-seq and ATAC-seq data to construct gene regulatory networks (GRNs), which are subsequently used for gene perturbation. They designed CO to account for the role of distal cis-regulatory elements, e.g. enhancers, as well as proximal promoters in the gene regulation system. For this purpose, they employed Cicero to determine the co-accessibility scores between peaks, provided by ATAC-seq data. These scores are then used to identify the interaction of distal regions with the target gene. Using CO, they have conducted multiple perturbation studies on different organisms and identified novel phenotypes resulting from transcriptional factor (TF) perturbation. In addition, they benchmarked CO’s performance using ChIP-seq data as ground truth against other state-of-the-art GRN methods across multiple mouse tissue samples. However, our evaluation reveals critical limitations in the implementation of their methodology, both in terms of ATAC-seq data integration as well as benchmarking. In this report, we first explain the limitations in their approach of integrating ATAC-seq data. We show that the proposed algorithm fails to account for distal regulatory interactions. After, we present the issues associated with their benchmarking algorithm and the data used for benchmarking. We show that their findings regarding the comparative performance of CO against other GRN inference methods is invalid and requires further evaluation. In conclusion, we detect multiple inaccuracies in this paper which undermine the validity of their published protocol and the results. The materials supporting our findings are accessible on GitHub^1^.

## 2 Integration of ATAC-seq data

In the published paper [1], in the section “*Identification of promoter and enhancer regions using scATAC-seq data*”, we read “*CellOracle uses genomic DNA sequence information to define candidate regulatory interactions. To achieve this, the genomic regions of promoters and enhancers first need to be designated, which we infer from ATAC-seq data. We designed CellOracle for use with scATAC-seq data to identify accessible promoters and enhancers (Extended Data Fig. 1a, left panel). Thus, scATAC-seq data for a specific tissue or cell type yield a base GRN representing a sample-specific TF-binding network*.”. In the next paragraph, we read “*To identify promoter and enhancer DNA regions within the scATAC-seq data, CellOracle first identifies proximal regulatory DNA elements by locating TSSs within the accessible ATAC-seq peaks. This annotation is performed using HOMER (http://homer.ucsd.edu/homer/). Next, the distal regulatory DNA elements are obtained using Cicero, a computational tool that identifies cis-regulatory DNA interactions on the basis of co-accessibility, as derived from ATAC-seq peak information. Using the default parameters of Cicero we identify pairs of peaks within 500 kb of each other and calculate a co-accessibility score. Using these scores as input, CellOracle then identifies distal cis-regulatory elements defined as pairs of peaks with a high co-accessibility score (≥0.8), with the peaks overlapping a TSS*.”

**Figure 1.**
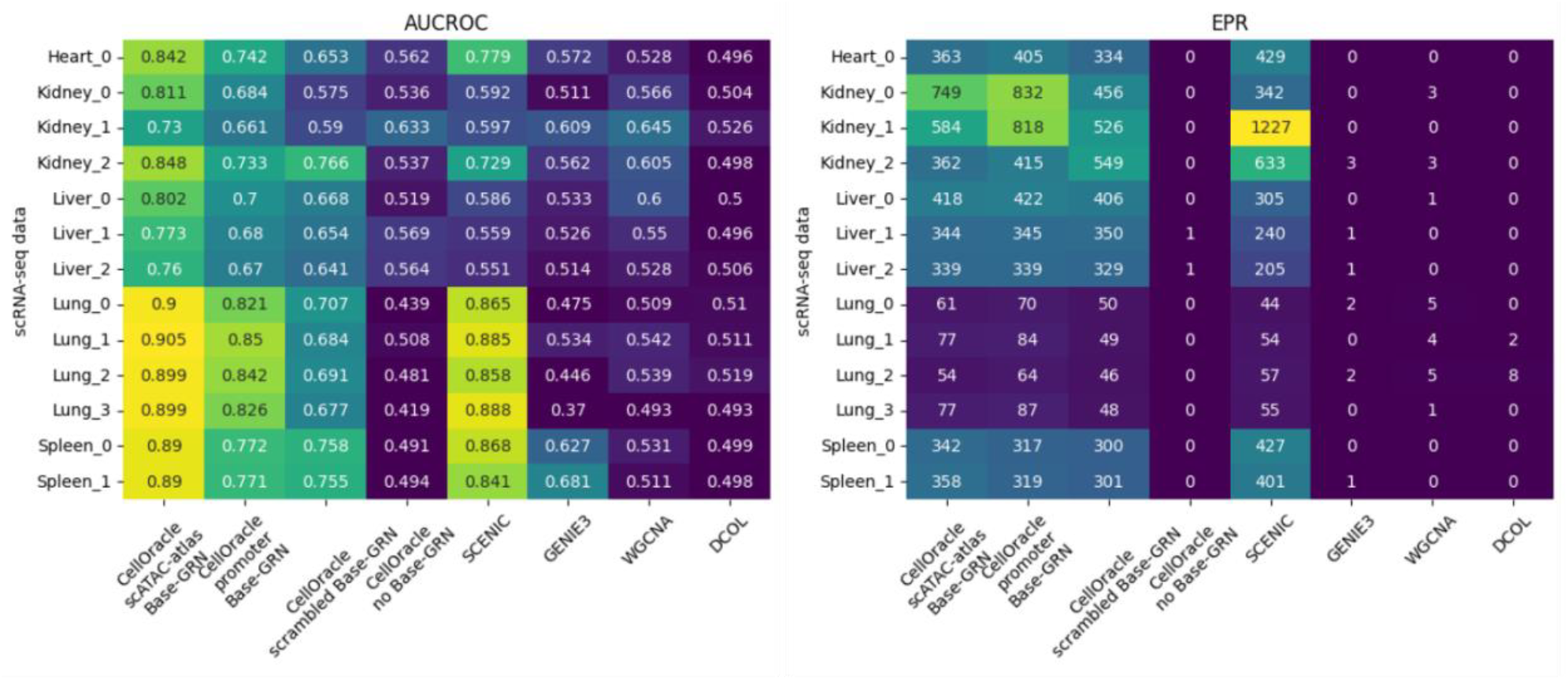
The results of the benchmarking process based on the original implementation

Here, we show that the pipeline of CO does not account for the distal cis-regulatory elements (enhancers) as claimed hereabove. CO first runs Cicero on the peak data to obtain the co-accessibility outputs in a [peak1, peak2, coaccess_score] format, which are continuous values within [-1, 1]. Then, similar to the published text, they take the following steps in the published tutoria^l2^ to obtain filtered peak-gene connections, which later will be used to derive the base GRN:

1. Function ***get_tss_info***: identify TSS peaks as the peaks that overlap the TSS of a gene. They have used a distance of 1000 base pair (bp) around TSS to find the intersection. This results in tss_annotated which is a dataframe with entries in the [peak, gene_name] format. This list contains potential proximal cis-regulatory elements, i.e. promotors, and their associated gene names. This creates around 17k peak-gene connections.
2. Function ***integrate_tss_peak_with_cicero:***
  a. Process Cicero co-accessibility data: replace *peak1* in Cicero connections with the corresponding gene name obtained in step 1. This gives *cicero_tss*, a dataframe with entries in the [peak_id, gene_name, coaccess_score] format.
  b. Filter *cicero_tss*: retain only positive scores. This reduces the non-zero connections resulting from Cicero from ∼210k to 110k.
  c. Merge TSS data and Cicero data:
    i. Assign co-accessibility scores of 1 to those peak-gene pairs present in *tss_annotated*.
    ii. Add these pairs to *cicero_tss* to obtain the complete list of peak-gene pairs.
3. Choose those peak-gene pairs with high co-accessibility scores to create a shortlist. They use a threshold of 0.8 on the co-accessibility scores for this purpose.

This pipeline in the official tutorial results in a total number of 15779 pairs of shortlisted peak-gene connections. We show that all 15779 connections belong to *tss_annotated* alone and no connection is originating from *cicero_tss*. This is because in step 2.c, assigning an artificial co-accessibility score of 1 to those peak-gene pairs in *tss_annotated* candidates these pairs as the top connections, where by the threshold of 0.8, no other peak-gene pairs remain in the shortlisted connections. As shown in step 1, *tss_annotated contains* the peaks intersecting TSSs and are independent of Cicero results. In other words, no distal cis-regulatory interactions are present in the network constructed by CO. This conflicts with the claims of CO to account for both proximal and distal cis-regulatory elements. In step 5, a more flexible threshold would include peak-gene pairs suggested by Cicero, as putative distal regulatory elements. However, we show that by choosing a more relaxed threshold of 0.5, still less than 1% of the final connections are taken from Cicero outputs. Thus, it is not clear how to choose this threshold to meaningfully integrate distal elements. Nonetheless, the choice of 0.8 as cutoff was also adopted by other studies using CellOracle, for example [2].

CO includes only those connections where given peaks either directly intersect with TSSs or are paired with peaks that intersect with TSSs (see steps 3 and 4). This creates very sparse peak-gene connections, with a median of 1 and a maximum of 6 peaks per gene. We also ran the pipeline of CellOracle on another multi-omics data with more peaks (the code is not provided as it is part of an ongoing study). We observed similar numbers; median of 1 and maximum of 8 peaks. For comparison, we also ran the pipeline of Scenic+ [2] and FigR [3] for the same data. To evaluate peak-gene connections, we define the domain of regulatory chromatin (DORC) scores as the number of peaks connected to genes [3]. We defined a threshold of 10 on peak connections to select DORC genes [3]. We obtained 62 DORCs for FigR, 4250 for Scenic+, and zero for CO.

## 3 Comparative benchmarking analysis of the GRN inference methods: algorithm, data, and results

For the first part, we received the data and Python code from the original authors, as these code and data were not officially released. The data encompasses inferred GRNs for various samples and tissues, along with the accompanying benchmarking code, which we made available online^3^.

### 3.1 Reproducing the results of the original paper

Using the code we received from the authors, we reproduced the reported results in Extended Data Fig. 2-A and B of the original paper [1]. To this end, we ran the code that receives the inferred GRN links for different methods as well as the ground truth as inputs and calculates AUROC and EPR scores, as shown in Figure 1.

**Figure 2.**
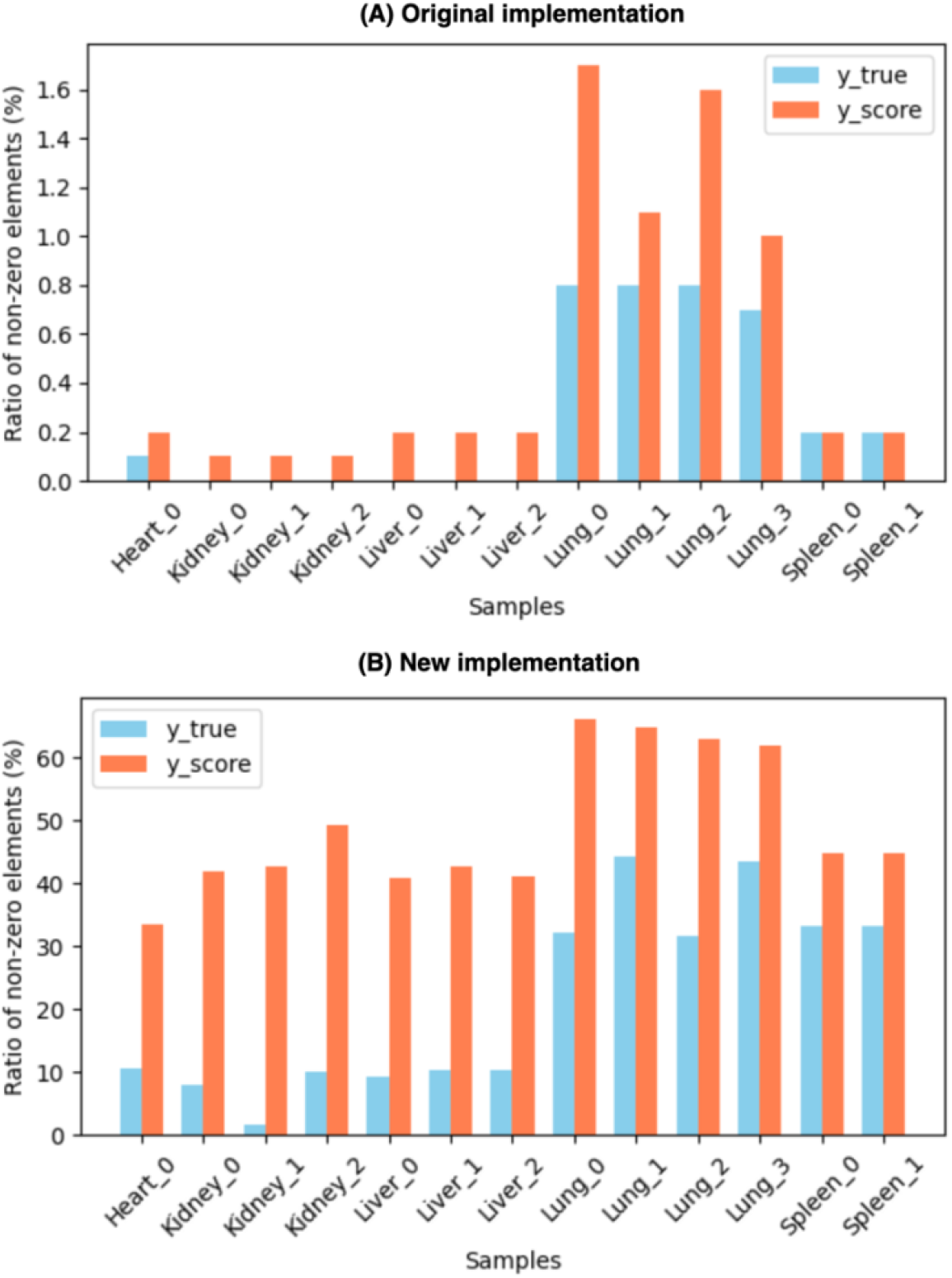
The ratio of non-zero elements obtained for *y_true* and *y_score* using (A) the original implementation and (B) the new implementation.

### 3.2 The original benchmarking algorithm and the associated issues

The authors calculated the AUROC and PR scores in two stages: first, by processing the inferred GRN and the ground truth to derive scores (*y_score*) and ground truth values (*y_true*)^4^; second, by using scikit-learn’s functions to compute the actual scores. In the first stage, the authors executed three steps to derive *y_score* and *y_true*^5^. (1) They created a comprehensive dataframe, *all_combinations*, containing all potential gene regulatory links. The aim of this step is to normalize comparisons between different GRN methods since methods such as CO and Scenic selectively filter out unlikely links using prior knowledge, while approaches such as GENIE3 do not, yielding a broader set of links. (2) They filtered this dataframe to retain only those links with transcription factors (TF) common to both the inferred GRN and the ground truth to ensure comparability. (3) Within *all_combinations*, they assigned a value of 1 to links present in the ground truth and 0 to those absent to create *y_true*. Similarly, they assigned the inferred values to the links in the inferred GRN and set 0 for the rest to derive *y_score*.

The methodology of the first two steps raises issues that need a closer look. The Python code for these steps is cited^6^ and is included below to support the ensuing argument.

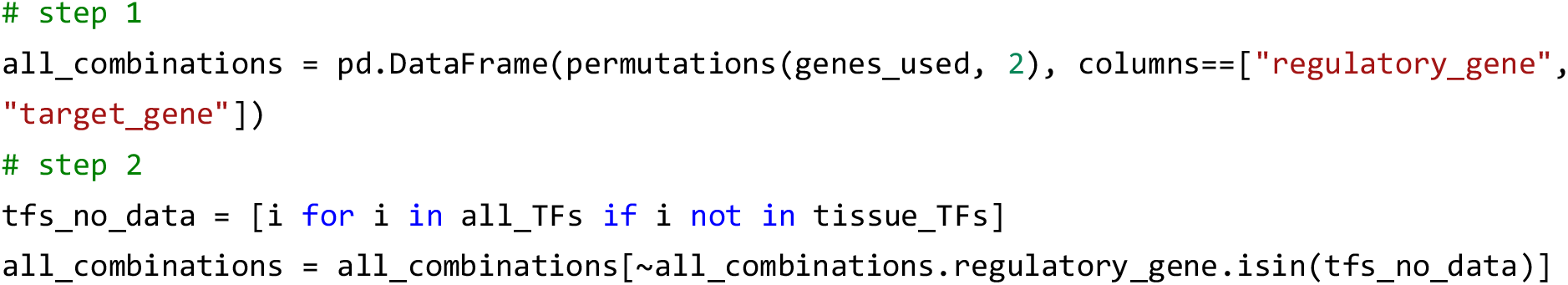

In this context, *genes_used* refers to the complete set of genes, encompassing both TFs and target genes. *All_TFs* denotes all TFs, while *tissue_TFs* specifies the subset of TFs present in the ground truth data for the particular tissue being evaluated. *regulatory_gene* is used synonymously with TF.

The *all_combinations* dataset, as constructed in the provided code, aggregates all potential gene interactions from the *genes_used* list. Of note, this step allows target genes, which are **not** listed in *all_TFs*, to be considered as regulatory genes. In step 2, the authors filtered out each link *(i, j)* in *all_combinations* when *i* is present in *all_TFs* but not in *tissue_TFs*. This step ensures that TFs are tissue specific. However, the method overlooks the links *(i, j)* where *i* is not a TF and therefore is certainly not present in the ground truth as TF either. Therefore, as those extra links are not filtered, they are included in the subsequent analysis and are assigned 0 in *y_true* and *y_score*. This leads to a marked imbalance between positive and negative values in calculating the scores. To illustrate the impact of this discrepancy, Figure 2-A displays the proportion of non-zero to zero elements, revealing that for most samples in the benchmarking, this ratio is below 0.2%. Such a pronounced imbalance between positive and negative instances could significantly compromise the reliability of the resulting scores.

### 3.3 Fixing the issues and recalculating benchmarking scores

To address the identified issue, we propose an alternative method for compiling *all_combinations*^*7*^. This method exclusively generates regulatory combinations where the regulatory gene is among the TFs confirmed in the ground truth data. This approach is crucial and also aligns with the authors’ original intention which was to only evaluate genes as regulatory genes if they are actual TFs. By recalculating *all_combinations* in this manner, we continue with the original methodology for defining *y_true* and *y_score*, to compute the scores. Using the new implementation, we see a notable improvement in the obtained ratio of non-zero elements in *y_true* and *y_score*, as given in Figure 2-B.

However, the ratio remains low in certain cases—for instance, a 2% ratio for *y_true* in the case of Kidney-1. This is attributed to the inherent sparsity in the connections from TFs to target genes.

Next, we obtain AUROC and EPR for CO and other GRN methods using the new implementation, as given in Figure 3. In addition, we calculated area under precision recall curve (AUPR) and F1 scores, as these two metrics are known to manage the class imbalance better than AUROC (see Figure 3). We can see that CO still performs comparatively well considering all metrics. However, in case of EPR, we see a significant drop in the scores for all methods. This indicates that the inferred GRN does not outperform the random models as significantly as reported in the original paper.

**Figure 3.**
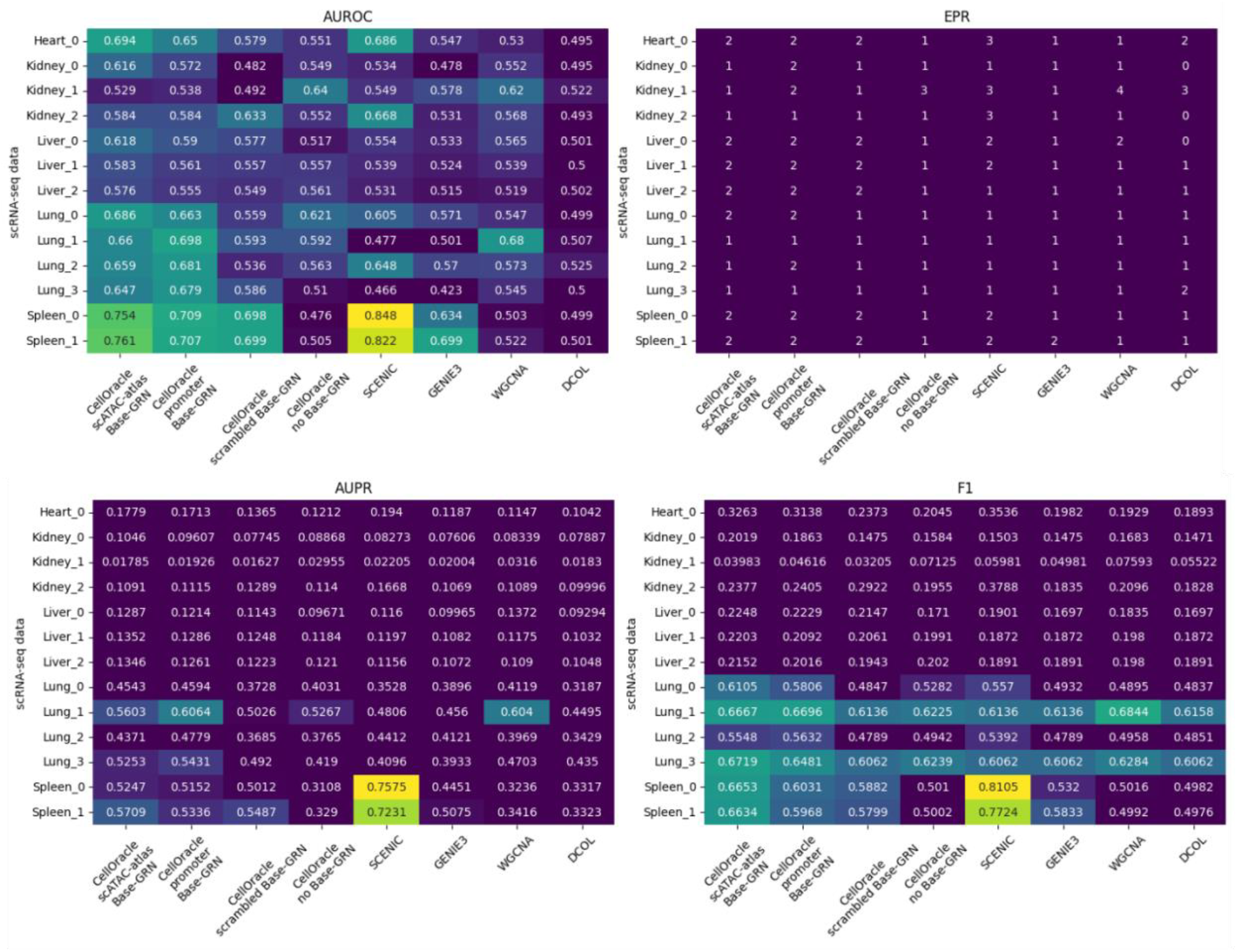
The results of benchmarking obtained after fixing the issues in the original implementation.

### 3.4 The coverage of the ground truth data

We reassessed the ground truth data employed for benchmarking in the original study, which claims to include 80 TFs (refer to the ‘Methods’ section titled “*Ground-truth data preparation for GRN benchmarking*”). However, our analysis revealed only 60 unique TFs across **all** tissues, where the number of actual TFs used to benchmark the inferred GRN for individual tissues goes as low as 2, as illustrated in Figure 4. Given that the inferred GRN could comprise over 1,000 TFs, such a low coverage of the ground truth, particularly for lung and spleen tissues, can significantly lower the validity of the comparative benchmarking scores. While the lack of comprehensive data is a common issue in GRN benchmarking, such a low coverage used in this study is rare. For instance, Sonawane [4] used benchmarking data with 19 to 59 TFs per tissue.

**Figure 4.**
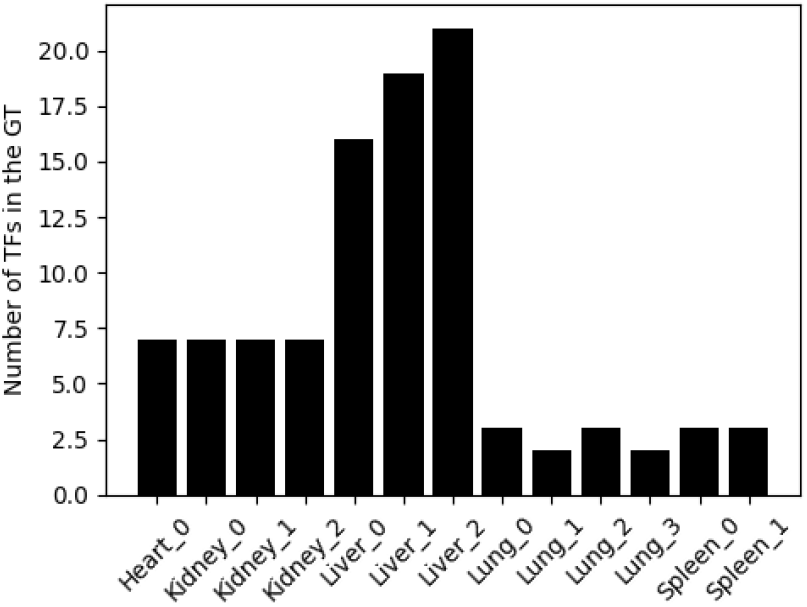
Number of TFs in the ground truth used for benchmarking.

## 4 Using gene expression data does not improve the accuracy of the GRN inference compared to the base GRN

Methods that employ prior knowledge, such as CO, use a base GRN derived from promoter priors or chromatin accessibility data, like ATAC-seq, to reduce false positives. This base GRN, expressed as binary connections, is refined by applying regression models to gene expression data. To highlight the base GRN’s contribution, the authors compared models using only regression (CellOracle-no Base GRN) against those incorporating a base GRN (such as CellOracle scATAC-atlas Base GRN), showing significant benchmarking score improvements with the latter. However, they did not quantify how gene expression data enhances GRN inference over the base GRN alone. Given the computational intensity of CO’s multiple regression models for each target gene, the added value of these models warrants investigation. We addressed this by shuffling the TF-target link values obtained from CellOracle scATAC-atlas base GRN, recalculating benchmarking scores, and comparing them to the unshuffled model via *t*-tests. The results suggest that the original model’s performance improvement over the shuffled GRN is minimal, as illustrated in Figure 5, lacking statistical significance (p-values of 0.4, 0.64, and 0.92 for AUROC, EPR, AUPR, and F1, respectively). This implies that the regression models utilized do not extract notable information from the expression data, thus failing to significantly enhance the regulatory insights beyond what is already indicated by chromatin accessibility data.

**Figure 5.**
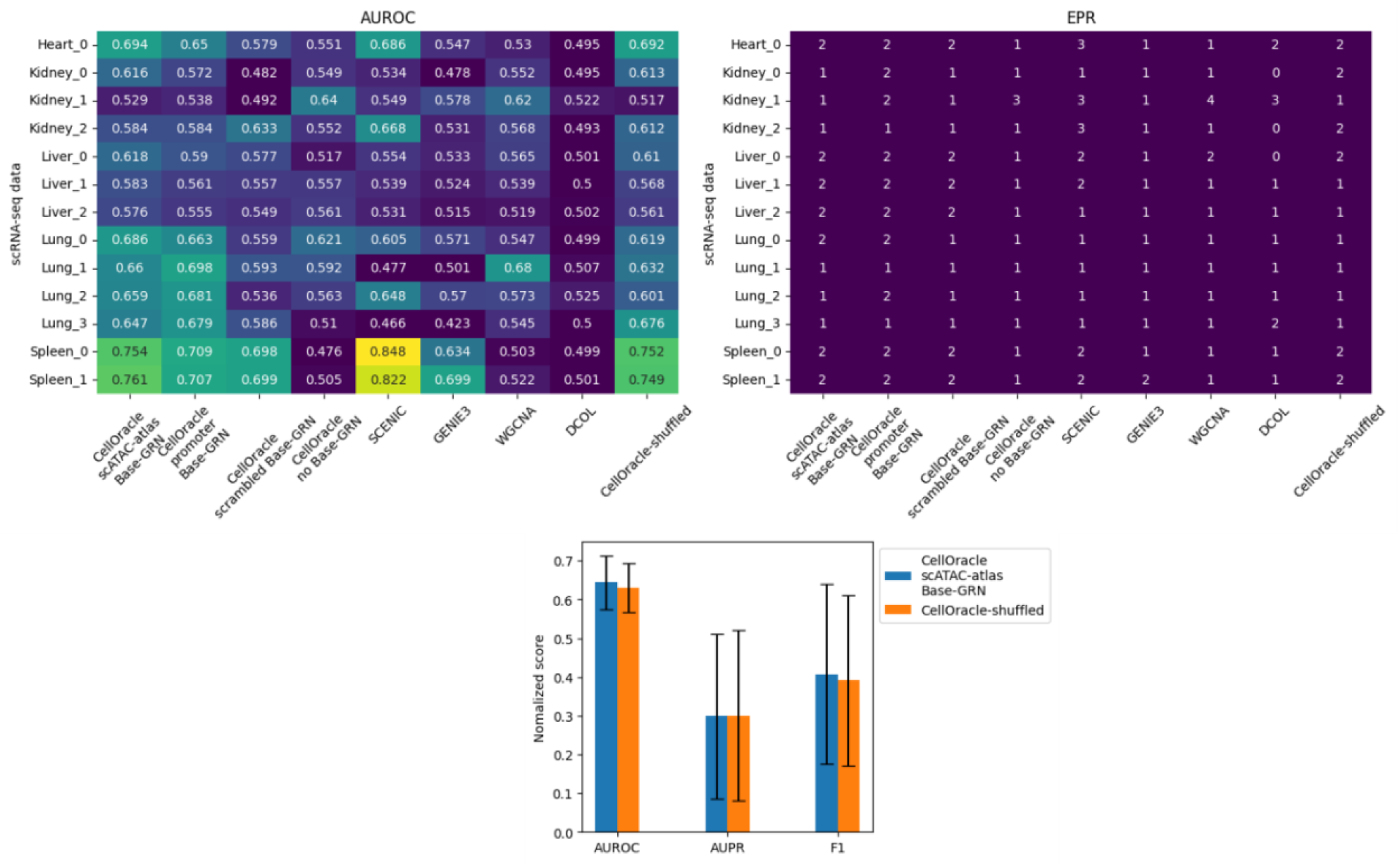
The benchmarking scores for the inferred GRN values versus randomly shuffled values.

## 5 Conclusion

This report uncovers several critical limitations within the study “*Dissecting cell identity via network inference and in silico gene perturbation*.” We showed that the proposed algorithm in CO fails to account for distal regulatory interactions. Also, we have pinpointed a substantial flaw in the original CO implementation that skews benchmarking scores and comparative analyses. Moreover, our evaluation reveals that the ground truth data used for benchmarking is not sufficient to allow for an effective comparison between different GRN inference methods. We also demonstrate that the regression model used in GRN inference does not significantly outperform the base GRN derived from ATAC-seq data. Given these findings, we advocate for further comprehensive re-evaluation of the method and results published in this paper to affirm the reliability of CO for GRN inference, a crucial step for accurate gene perturbation simulation.

## Footnotes

1 https://github.com/janursa/CO_evaluation

2 https://github.com/morris-lab/CellOracle/blame/b28fdbfdfbd9c55a80b1471d88a59f84ea30d662/docs/notebooks/01_ATAC-seq_data_processing/option1_scATAC-seq_data_analysis_with_cicero/02_preprocess_peak_data.ipynb#L1

3 https://github.com/janursa/CO_evaluation

4 https://github.com/janursa/CO_evaluation/blame/2f385882e4476dc5cf03810b46692c7eac467ccd/CO_evaluation/utils/CO_benchmark.py#L214

5 https://github.com/janursa/CO_evaluation/blame/2f385882e4476dc5cf03810b46692c7eac467ccd/CO_evaluation/utils/CO_benchmark.py#L271

6 https://github.com/janursa/CO_evaluation/blob/8e265b9ce191852bf36f4faa53a4d728838850bf/original_code/GRN_benchmarking.py#L292

7 https://github.com/janursa/CO_evaluation/blame/d94f18eb7b7c7e08b3eb2287a010e2c88a0437bc/CO_evaluation/utils/CO_benchmark.py#L238

